# VarGenius-HZD allows accurate detection of rare homozygous or hemizygous deletions in targeted sequencing leveraging breadth of coverage

**DOI:** 10.1101/2021.06.21.449209

**Authors:** Francesco Musacchia, Marianthi Karali, Annalaura Torella, Steve Laurie, Valeria Policastro, Mariateresa Pizzo, Sergi Beltran, Giorgio Casari, Vincenzo Nigro, Sandro Banfi

## Abstract

**Motivation:** Homozygous deletions (HDs) may be the cause of rare diseases and cancer and their discovery in targeted sequencing is a challenging task. Different tools have been developed to disentangle HD discovery but a sensitive caller is still lacking.

**Results:** We present VarGenius-HZD, a sensitive and scalable algorithm that leverages breadth-of-coverage for the detection of rare homozygous and hemizygous single-exon deletions (HDs). To assess its effectiveness we detected both real and synthetic rare HDs in fifty exomes from the 1000 Genomes Project obtaining higher sensitivity in comparison with state-of-the-art algorithms which missed at least one event each. We then applied our tool on targeted sequencing data from patients with Inherited Retinal Dystrophies and solved five cases that still lacked a genetic diagnosis.

**Availability and implementation:** We provide VarGenius-HZD either stand-alone or integrated within our recently developed software enabling the automated selection of samples using the internal database. Hence, it could be extremely useful for both diagnostic and research purposes. Our tool is available under GNU General Public License, version 3 at: https://github.com/frankMusacchia/VarGenius-HZD

Supplementary information is available online.

## INTRODUCTION

Next generation sequencing (NGS) is commonly used to unveil genetic causes of diseases and Whole Exome Sequencing (WES) has become one of the most commonly used diagnostic tools both in the clinic and in several programs investigating rare genetic diseases. Rare diseases collectively affect a significant fraction of the population (estimated to be about 4-5%) (Boycott et al., 2013; Wright et al., 2018) with a resulting strong impact on health-care costs and mortality rates. Currently, the standard protocol to investigate rare diseases includes multiple clinical diagnostics assays. Nonetheless, half of the cases still remain without a diagnosis (Demos et al., 2019; Hartley et al., 2020; Shashi et al., 2014). One of the reasons for this is the limited knowledge of how to detect Copy Number Variation (CNV) from sequencing data, when, it is estimated that about 12% of the genome in the human population is subject to copy number changes (Redon et al., 2006; Yuan et al., 2020). To detect CNVs, diagnostic laboratories often use multiplex ligation-dependent probe amplification (MLPA) and array comparative genomics hybridization analysis (ArrayCGH) prior to executing NGS-based analysis (Vasson et al., 2013). Yet, both methods have high ranges in resolution (from kilobases to megabases) and add complexity to the overall patient screening process. Whole Genome Sequencing (WGS) data are more even in coverage in comparison to WES making it more reliable for CNV calls. However, due to massive use of WES in diagnostics, there is a need for reliable methods to infer CNVs from exome data as well (GR et al., 2016; Lelieveld et al., 2016; Stark et al., 2016). Indeed, leveraging the sequencing outcome to detect CNVs offers potential advantages leading to increased diagnostic yield without increasing laboratory costs (Lelieveld et al., 2016; Pfundt et al., 2017).

Several CNV-detection algorithms for WES data have been developed, all of which rely on the use of depth-of-coverage (DoC) from multiple samples to infer copy numbers (Fromer & Purcell, 2014; Krumm et al., 2012; J. Li et al., 2012; Plagnol et al., 2012). Unfortunately, the CNV search is hampered by biases due to differences in capture protocol efficiency, the presence of GC-rich regions and different coverage resolutions that influence DoC, among others (Auer et al., 2016; Guo et al., 2013; Hong et al., 2016). Such heterogeneity complicates the downstream analysis of the detected events leading to false positives (Feng et al., 2015; Guo et al., 2013; P. S. Samarakoon et al., 2016; Tan et al., 2014) while compromising the ability to reliably detect CNVs when these span less than three exons (Auton et al., 2015; Feng et al., 2015b; Hong et al., 2016; Lelieveld et al., 2016; Moreno-Cabrera et al., 2020). Even though CNV detection could represent a valuable complementary way to analyze NGS data, the low concordance of detected events suggests that the algorithms designed so far are yet to be optimised (Hong et al., 2016; Moreno-Cabrera et al., 2020; Tan et al., 2014; Yao et al., 2017). Moreover, comparative works demonstrated that these results are often difficult to replicate despite the high specificity and sensitivity declared (Sadedin et al., 2018). One method to overcome these issues could be to generate a consensus of variants called by different algorithms (Moreno-Cabrera et al., 2020). However, to use any of these approaches, the user needs to prepare BAM files for unrelated samples sequenced with the same target writing ad-hoc scripts making such analyses difficult for those laboratories that do not have bioinformatics expertise. Therefore, the implementation of a fully automated CNV workflow along with different methods to investigate CNVs in WES data beyond the DoC strategies is of high importance for the scientific community.

Single-exon homozygous/hemizygous deletion (HD) detection methods, which compare normalized coverage values among samples produced with the same kits already exist (e.g. Atlas-CNV, CoNVaDING, DECoN and HMZDelFinder) (Chiang et al., 2019; Fowler et al., 2016; Gambin et al., 2017; Johansson et al., 2016). While Atlas-CNV and CoNVaDING, as suggested by the authors, can only be used with high coverage sequencing data (e.g. small targeted gene panels), HMZDelFinder and DECoN are ad-hoc tools for exonic CNV detection. However, these tools are based on the assumption that the data has a defined distribution and hence require intra- and inter-samples homogeneity (Sadedin et al., 2018).

To overcome these challenges, we developed a new algorithm for the detection of rare single-exon HDs which exploits breadth-of-coverage (BoC) and we named it VarGenius-HZD (where HZD stands for homozygous/hemizygous deletion detection). Additionally, we automated its execution along with that of ExomeDepth and XHMM within our recently developed software that we devised for variant detection analysis and management of samples, i.e., VarGenius (Musacchia et al., 2018). This software is now able to automatically pick selected samples generated with the same target and to perform CNV calling separately on autosomes and sex chromosomes and in parallel across different cores of a High Performance Computing (HPC) system managed with Portable Batch System (PBS) scheduler. VarGenius-HZD algorithm is either integrated within VarGenius software where it scales across HPC nodes, or is available in a stand-alone version which takes as input a list of manually selected BAM files and allows scaling across CPU cores.

We have validated our algorithm using 50 samples from the 1000 Genomes Project (1KGP) (https://www.internationalgenome.org/) in which we detected both existing and artificially inserted HDs. For these test cases we compared VarGenius-HZD results with those of HMZDelFinder, DECoN and ExomeDepth and our algorithm obtained the highest sensitivity. Furthermore, we applied VarGenius-HZD on targeted sequencing data from a cohort of 188 individuals with Inherited Retinal Dystrophies (IRDs) resolving 5 out of 64 undiagnosed cases by identifying pathogenic HDs which were then experimentally validated.

## SYSTEM AND METHODS

### NGS procedures

The 188 subjects considered in this study were selected for targeted sequencing after being assessed at the Referral Centre for Inherited Retinal Dystrophies of the Eye Clinic at Università degli Studi della Campania ‘Luigi Vanvitelli’ (VRCIRD) (Table 1). Peripheral blood samples were collected upon written informed consent of the patient or their parents/legal guardians (for minors). All procedures adhered to the tenets of the Declaration of Helsinki, and were approved by the Ethics Board of Fondazione Telethon and Università degli Studi della Campania ‘Luigi Vanvitelli’.

**Table 1.** Summary of samples used from the VRCIRD cohort.

| platform | CREv1 | CCP | ID | Total | Solved Cases | Examined |
| --- | --- | --- | --- | --- | --- | --- |
| NextSeq500 | 14 | 51 | 123 | 188 | 124 | 64 |

DNA samples from peripheral blood were processed using the Illumina NextSeq500 (Illumina inc., San Diego, CA, USA). Three different enrichment kits (Agilent Technologies) were used for library preparation. In particular, 123 samples were sequenced using the Agilent ClearSeq Inherited Disease (ID), 51 samples using the SureSelect Clinical Constitutional Panel (CCP) and 14 samples were prepared using the SureSelect Clinical Research Exome version 1 (CREv1) (Table 1). BCL files were processed using Illumina bcl2fastq. Raw fastq files were processed using our previously developed software (Musacchia et al., 2018).

#### Calling homozygous deletions leveraging BoC

The VarGenius-HZD algorithm is written in PERL and R programming languages and needs the execution of three steps: sample selection, pre-processing and rare HD detection (Figure 1). The *samples selection* step is automated within the VarGenius software by querying the PostgreSQL database for unrelated samples sequenced with the same target while for the stand-alone version this step is manual i.e. the user must provide a file with the paths to the BAM files and the BED file for the target sequenced.

**Figure 1:**
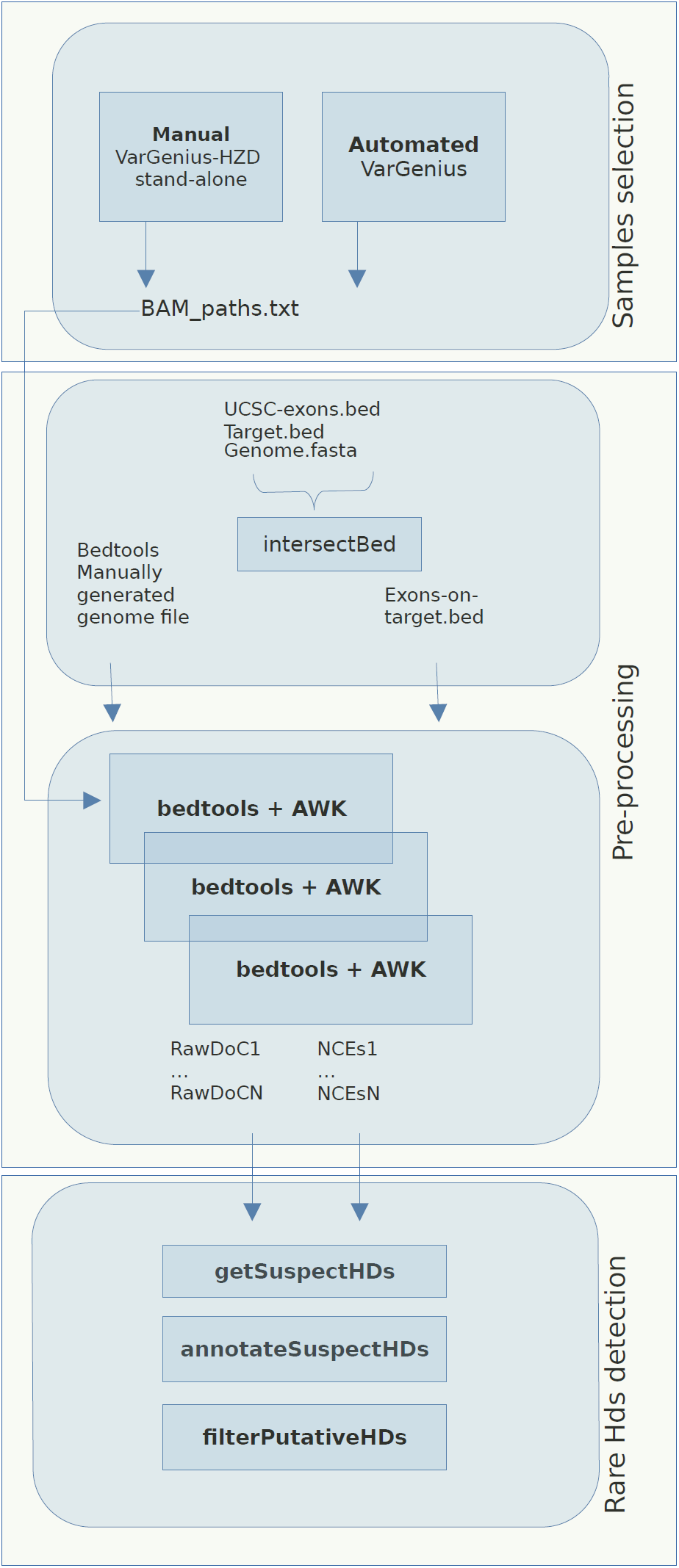
VarGenius-HZD workflow. The workflow of our algorithm consists of three steps: 1. samples selection, which is automated in VarGenius software and manual in the stand-alone version; 2. the pre-processing, which includes the generation of NCEs files and raw DoC information; 3. the rare HDs detection step which involves the calculation of NCE frequencies, the detection of putative HDs and the annotation of such regions for variant prioritization.

The *pre-processing* step aims to generate an exons-on-target intervals file. To this end, BED files for genes and exons were downloaded from the University of California Santa Cruz (UCSC) platform (https://genome.ucsc.edu/cgi-bin/hgTables) and included within the package. We selected UCSC genes as track, Hg19 as genome assembly, start and end of exons/genes and BED as output format. The BED file is then intersected with the target file using bedtools intersect. This procedure is executed only once per-target.

Furthermore, each BAM file undergoes bedtools *coverage* to compute the BoC and the DoC of the previously generated exonic intervals using this command: **bedtools coverage -a input.bam -b exons_on_target.bed -g ucsc.hg19.genomefile -sorted**. As suggested in the bedtools guidelines, we used the -sorted parameter and a *genome file* in input to accelerate such computation. The *genome file* was generated with the following commands: **samtools faidx ucsc.hg19.fa; cut -f1,2 ucsc.hg19.fa.fai > ucsc.hg19.genomefile** (https://bedtools.readthedocs.io/en/latest/content/tools/coverage.html). The resulting output, containing the BoC, is filtered to select only exons with BoC<0.2 (<20% of the exon covered) and annotated with the UCSC genes for downstream analyses. This procedure generated two tab separated files (TSV) for each sample: one containing putative non-covered exons (NCEs) and another containing raw DoC for the exons-on-target.

The third step aims at *rare HDs detection* using the two TSV files previously produced. First, all NCEs files are loaded within a unique array. Second, putative rare HDs are obtained by computing their frequency and selecting those where it is lower or equal to 2 (this parameter can be customized). Third, the exonic raw DoC of parents (whenever available), proband and average across all samples are added. Once complete, the annotated table of putative rare HDs is provided and manual filtering of relevant calls based on the difference in coverage between the proband and her/his parents and between the proband and the overall dataset can be performed downstream (Figure 2).

**Figure 2:**
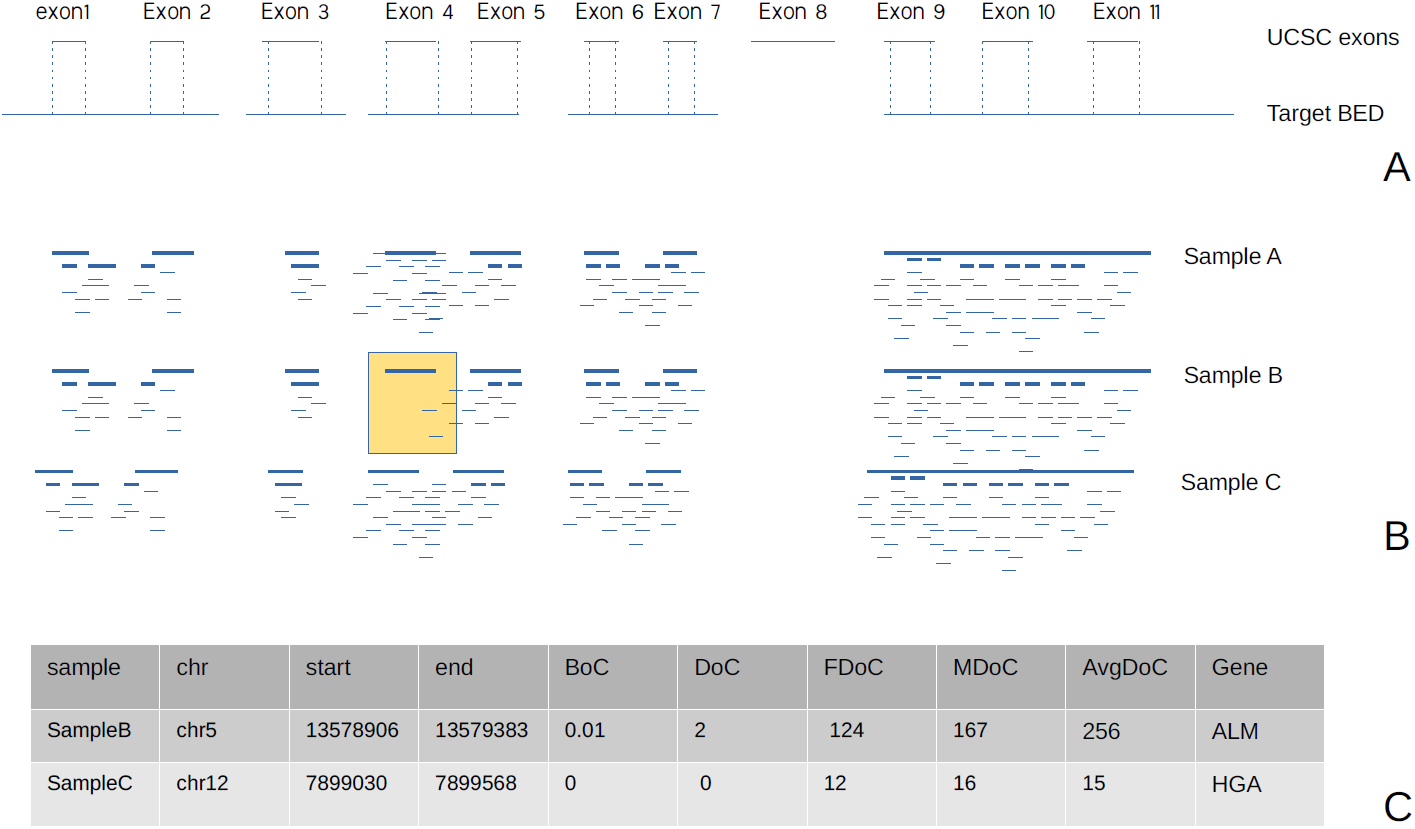
VarGenius-HZD algorithm and results ilustration. VarGenius-HZD leverages BoC along with DoC which is used as follows: (A) the target BED used for the sequencing is intersected with UCSC exon intervals to obtain an exon-on-target file which is used to compute the BoC and DoC exploiting bedtools coverage. (B) NCEs for each sample are counted and only those with frequency lower or equal to 2 are retained as putative HDs (e.g. exon 4 in figure B). (C) the tabular output contains statistics for putative HDs: chromosome, start and end, the BoC for the subject sample, the DoC for the parents (FDoC and MDoC) and average exon DoC for the overall dataset.

#### 1KGP WES dataset

We selected 50 samples for which genome wide CNV calls were available and from the 1KGP data as in (Gambin et al., 2017) from (http://ftp.1000genomes.ebi.ac.uk/vol1/ftp/phase1/data/) and their consensus target BED file from (http://ftp.1000genomes.ebi.ac.uk/vol1/ftp/phase1/analysis_results/supporting/exome_pull_down/20120518.analysis_exome_targets.consensus.annotation.bed). Variant Calling Format (VCF) files with genotypes were downloaded from (http://hgdownload.cse.ucsc.edu/gbdb/hg19/1000Genomes/). First the VCF file was intersected with the consensus BED files using bedtools intersect. Then, we filtered the VCF for rare HDs selecting those calls with *sv_type=del* within the VCF *info* field and where only one sample had *gt=1|1*. BAM files for the 50 samples were used with ExomeDepth, HMZDelFinder, DECoN and VarGenius-HZD. Results from the four tools are in Supplementary tables 2, 3, 4 and 5. HDs called were further filtered with the following approaches: for ExomeDepth and DECoN we retained only the calls with reads.ratio<0.001; HMZDelFinder provided a filtered output, hence we did not apply any filter; VarGenius-HZD produced a filtered output as well and we have picked those calls where the average DoC was greater than 50.

#### Synthetic deletion detection

The 50 samples from 1KGP were also used to conduct a test using simulated deletions generated with bedtools. We inserted 5 HDs in 5 distinct samples (NA06989, NA07347, NA12058, NA12748, NA12830) choosing regions that we have visually inspected and had sufficient coverage surrounding the chosen HDs (>20x) across the overall dataset (Table 2). The commands used to generate such deletions were: **bedtools intersect -a sample.bam -b deletion_i.bed -v > sample_deleted.bam; samtools sort sample_deleted.bam > sample_deleted_sort.bam; samtools index sample_deleted_sort.bam**.

**Table 2.** The 5 simulated HDs inserted in samples of 1KGP dataset.

| sample | chr | start | end |
| --- | --- | --- | --- |
| NA06989 | 21 | 48063447 | 48063551 |
| NA07347 | 21 | 27326904 | 27327003 |
| NA12058 | 21 | 35091133 | 35091161 |
| NA12748 | 21 | 10906904 | 10907040 |
| NA12830 | 21 | 40188932 | 40189015 |

HDs were filtered with the following methods only for the 5 samples: DECoN and ExomeDepth: reads.ratio<=0.001. VarGenius-HZD: raw average DoC >50; HMZDelFinder: no filtering.

#### Recall, precision and specificity scores

To compare results from different tools we calculated recall, precision and specificity scores with the following formula: Recall= TP/(TP+FP); Specificity= TN/(TN+FP); Precision: TP/(TP+FP). In this formula true positives (TP): are HDs called and present in the 1KGP VCF; true negatives (TN): HDs not called that are not present; false positives (FP): HDs called that are not present; false negatives (FN): HDs not called that are instead in the 1KGP VCF.

### Analysis of the VRCIRD cohort

#### Automated CNV detection workflow

All samples of the VRCIRD cohort were subject to SNVs/Indels and CNV calling. We used the GATK3.8 BestPractices with default parameters (Van der Auwera et al., 2013). Alignment was using BWA (H. Li & Durbin, 2010), PCR duplicates were marked with PICARD MarkDuplicates (Broad Institute, n.d.), and, further BAM pre-processing was performed using BaseRecalibrator prior to variant calling with HaplotypeCaller. We performed GATK *hard filtering* with VariantFiltration and Annovar was used for the annotation (Wang et al., 2010). Further parsing of the VCF file and annotation table was performed to provide an XLS tabular output to the physician. Sample information (such as sample and analysis name, gender, kinship and target used) provided within the samples sheet were parsed and stored within the PostgreSQL database.

CNV detection was performed with XHMM, ExomeDepth and VarGenius-HZD, as follows: the software executed a query to the PostgreSQL database to obtain samples which are sequenced with the same target. VarGenius automatically picked the BAM files to provide as input for the tools from its results folders using sample identifiers, kinship, gender and the target used. XHMM and ExomeDepth were executed following the author’s guidelines and with default parameters. Autosomes and sex chromosomes were analyzed separately to avoid gender biases. Once finished, all the CNVs called (herein “calls”) were annotated using AnnotSV (V et al., 2018) (Figure 3). Causative SNVs/Indels for all subjects were investigated and a subset of 124 cases received a diagnosis. The remaining 64 cases were subject to CNV prioritization. Yet, in this work, we specifically discuss only HDs.

**Figure 3:**
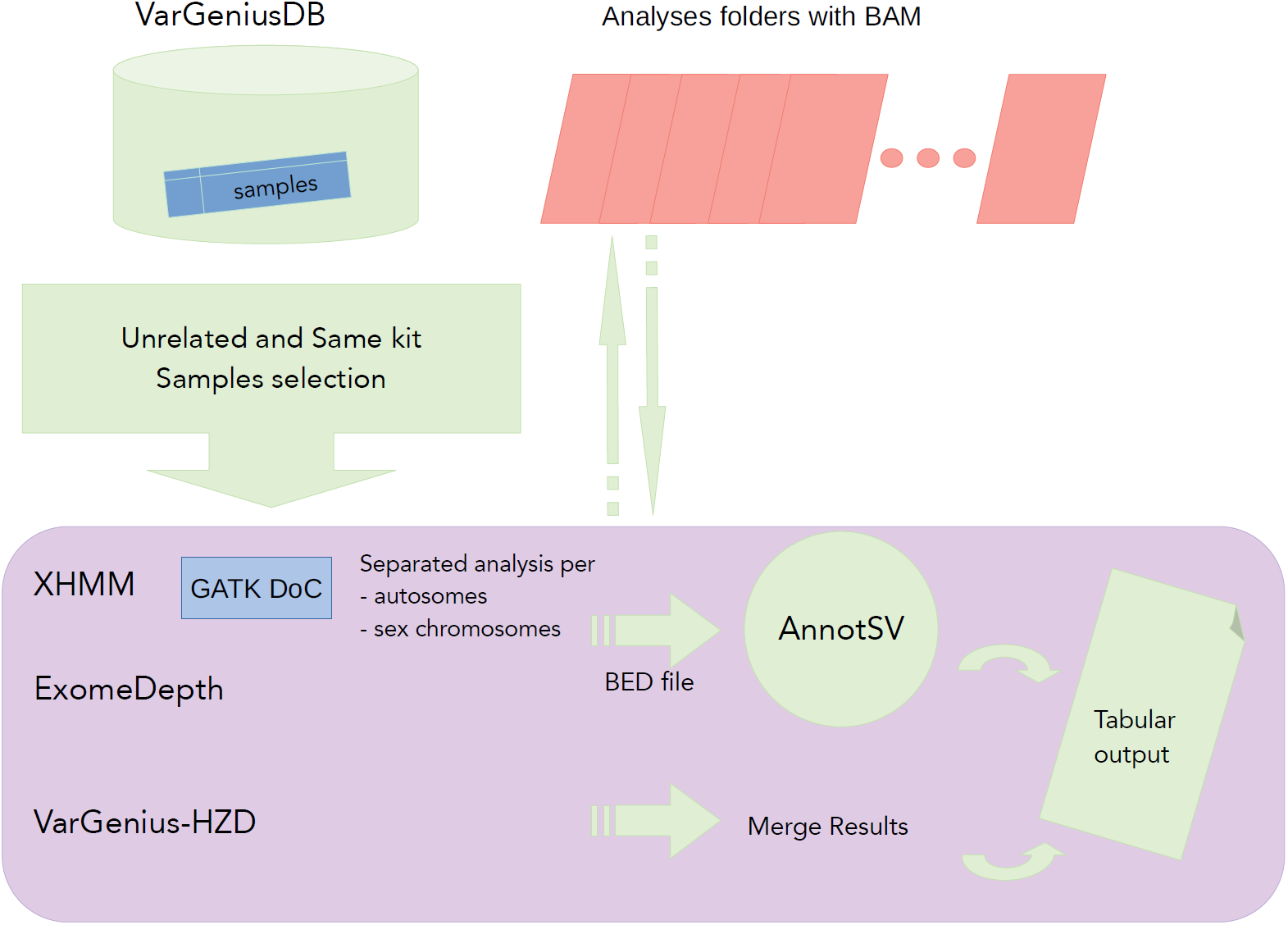
Flowchart of CNV detection and annotation pipeline in VarGenius. This is performed using XHMM, ExomeDepth and VarGenius-HZD algorithm. Several unrelated samples must be used for such analyses thus VarGenius collects samples identifiers from the database querying for samples sequenced with the same target and considering the kinship. XHMM requires the use of GATK DepthOfCoverage with specific parameters. This is called for all samples parallelizing the execution within the cluster. Once all tools produced their calls, results are merged within a unique tabular output and are annotated using AnnotSV.

#### HDs filtering for patients

Manual inspection in Integrative Genomics Viewer (IGV) of detected HDs was performed for 64 undiagnosed cases which could not be solved with a causative SNV/Indel (Table 1). To increase the probability of diagnosis through known disease genes, we filtered the resulting calls using our internal panel of known retinopathy genes (data not published). Selection of resulting events to inspect in IGV as a first tier validation, was manually performed keeping into account AnnotSV annotation (OMIM, Decipher) and different scores depending on the tool. For ExomeDepth we considered the Bayes factor (BF>10); for XHMM we used the mean normalized DoC (MEAN_RD); for VarGenius-HZD we selected calls with average raw DoC > 30. BAM files for the proband and her/his parents (whenever available) or for different probands were loaded as controls.

## RESULTS

### BoC can be used along with DoC to detect rare HDs

#### Results comparison with 1KGP data

To compare the performances of different algorithms in detecting rare HDs, we applied VarGenius-HZD, ExomeDepth, HMZDelFinder and DECoN to 50 samples from the 1KGP selecting only rare HDs (see Methods). The resulting calls are in Supplementary Table 1. One of the five HDs found in the 1KGP VCF file appeared to be a false positive (sample NA19473 - position: chr9:107366951) when inspected in IGV as it was covered by ~60 reads and none of the tools detected it (Supplementary Table 1 and Supplementary Figure 1). Therefore, we considered only 4 real HDs. ExomeDepth detected only 1 event while 2 were filtered out (Supplementary Table 1 and Supplementary Table 3); HMZDelFinder found only one HD (Supplementary Table 1 and Supplementary Table 4); DECoN could detect only 1 HD and 1 was filtered out (Supplementary Table 5). VarGenius-HZD was able to detect the highest number of true positive events: 3 out of 4 and one was filtered out (Supplementary Table 1 and Supplementary Table 2). The HD in sample NA11919 (position: chr5:140222138) was called by all tools. For the sake of curiosity, we inspected in IGV the regions near single nucleotide homozygous variants present in total in four samples: NA20798, NA19137, NA18504, NA18950 (Supplementary Table 6). Intriguingly, in sample NA20798, we found that the genes *CFHR3* and *CFHR1* were deleted. We could infer the call after inspection of the coverage of the nearby *CFHR* and *CFHR4* genes coverage and through comparison with control samples (Supplementary figure 2). This event was correctly detected by all tools but was not included in the 1KGP results (Supplementary Tables 1-5). Furthermore, a putative HD of gene *UGT2B28* in sample NA18504 was detected by VarGenius-HZD, ExomeDepth and filtered out by DECoN and visually confirmed by comparing samples coverage in IGV (Supplementary figure 3 and Supplementary Tables 2-5).

To assess our results we computed precision, recall and specificity scores for all tools (none of the newly discovered variants was included in such calculations). VarGenius-HZD obtained higher recall, specificity and precision when compared with ExomeDepth and DECoN (~2-5% vs ~0.4%). However, VarGenius-HZD results are comparable with those of HMZDelFinder: we obtained higher recall with our tool (75% vs 25%), but higher specificity (10% vs 2%) and precision (10% vs 6%) with HMZDelFinder (Table 3). HMZDelFinder appeared to be the most precise tool returning very few events to inspect reducing the number of FPs but at a cost of missing TP events and losing in sensitivity. The highest number of true positive calls was instead obtained by VarGenius-HZD. All tools found one additional putative TP HD. Yet, this variant should be experimentally confirmed which was out of the scope of this study.

**Table 3.** Precision/Recall/Specificity obtained by the tools with the 50 samples from 1KGP dataset.

| Algorithm | TotalPutativeHZDel | TP | TN | FN | FP | recall | specificity | precision | NewTP |
| --- | --- | --- | --- | --- | --- | --- | --- | --- | --- |
| ExomeDepth | 274 | 1 | 1 | 3 | 273 | 0,25 | 0,0036 | 0,0036 | 2 |
| VarGenius-HZD | 51 | 3 | 1 | 1 | 48 | 0,75 | 0,0204 | 0,0588 | 2 |
| HMZDelFinder | 10 | 1 | 1 | 3 | 9 | 0,25 | 0,10 | 0,10 | 1 |
| DECoN | 267 | 1 | 1 | 3 | 266 | 0,25 | 0,0037 | 0,0037 | 1 |

#### Detection of synthetic HDs

To evaluate our results with synthetic data, we simulated five deletions in five distinct samples of 1KGP selected randomly from our cohort of fifty. After running the tools using the fifty samples, to ameliorate positive and negative identification, we considered only HD calls for the selected samples after filtering as described in Material and Methods. ExomeDepth and DECoN could not find any of the simulated events, HMZDelFinder missed only one and VarGenius-HZD found them all (Table 4). Since all the detected events were true positives, HMZDelFinder obtained 100% precision and 80% recall. On the contrary, VarGenius-HZD detected all the true positive synthetic deletions inserted as well as one additional putative false positive, reaching 100% recall and 83% precision. We speculate that downstream CNV filtering is always needed and performed in several ways (e.g. visual inspection in IGV, gene panel selection, clinical phenotype, etc.) hence, for clinical diagnostics, a higher recall at the cost of a bit of downstream manual work would be preferable as it leads to a higher number of positive genetic diagnoses.

**Table 4.** Precision/recall/specificity obtained by the tools used with the synthetic HDs test.

| Algorithm | TotalCalls | TotalFiltered | TP | TN | FP | FN | recall | specificity | precision |
| --- | --- | --- | --- | --- | --- | --- | --- | --- | --- |
| HMZDelFinder | NA | 4 | 4 | 0 | 0 | 1 | 0.8 | 0 | 1 |
| VarGenius-HZD | 4201 | 6 | 5 | 0 | 1 | 0 | 1 | 0 | 0.83 |
| DECoN | 38234 | 45 | 0 | 0 | 45 | 5 | 0 | 0 | 0 |
| ExomeDepth | 3949 | 3 | 0 | 0 | 3 | 5 | 0 | 0 | 0 |

### Automated SNV/Indel and CNV calling

We had previously developed VarGenius to execute SNV/Indel calling and annotation exploiting the GATK BestPractices pipeline. VarGenius is able to scale across nodes of an HPC cluster with PBS scheduling system and to construct a PostgreSQL database to store sample information enabling several queries for general genetics investigation.

We have embedded within VarGenius the execution of ExomeDepth, XHMM and VarGenius-HZD for CNV analysis. Validation of results from ExomeDepth demonstrated high specificity and sensitivity for the detection of rare variants (Ellingford et al., 2017; Plagnol et al., 2012; P. Samarakoon et al., 2014) while further studies suggested Conifer and XHMM for the low occurrence of false-positives, but with the disadvantage of a low detection sensitivity (Fromer & Purcell, 2014; Guo et al., 2013; Krumm et al., 2012; P. S. Samarakoon et al., 2016; Yao et al., 2017). Thus, XHMM, Conifer and ExomeDepth are the tools best adapted to detect rare variants (Zhao et al., 2013). However, Conifer has not been updated for years and it relies on the installation of old versions of R rendering difficult its integration within an automated pipeline.

State-of-the-art CNV detection tools need as input several BAM files sequenced with the same enrichment kit and different analyses should be performed for autosomes and sex chromosomes to avoid ploidy biases. We automated this process in VarGenius by querying for their numeric identifiers within the PostgreSQL database (see Methods section and Figure 1). Several software are available to execute scalable CNV analysis for targeted sequencing data such as bcbio (https://github.com/bcbio/bcbio-nextgen) and nf-Sarek (Garcia et al., 2020), yet they require the user to manually select the BAM files to use. Other open source tools (e.g. Hpexome, HemoMIPs and Swift/T) allow automated and scalable detection of SNV/Indel b using multiple samples, but not CNVs (Table 5). Since VarGenius automates the complete workflow needed to execute CNV analysis, it can be a valuable resource for laboratories lacking bioinformatic expertise.

**Table 5.** Availability of SNV/CNV analysis automation in existing open source software.

| Software | SNV/Indel Calling | CNV calling | scalability | Automated dataset creation |
| --- | --- | --- | --- | --- |
| bcbio | yes | yes | yes | no |
| Nf-sarek | yes | yes | yes | no |
| Hpexome | yes | no | yes | no |
| HemoMIPs | yes | no | yes | no |
| Swift/T | yes | no | yes | no |

We have applied ExomeDepth, XHMM, VarGenius-HZD and HMZDelFinder with default parameters to the 188 samples of the IRD cohort running different analyses for different enrichment kits. We first filtered only calls for the 64 unsolved cases using our panel of retinopathy genes and the thresholds suggested in the corresponding user guides (see Methods). ExomeDepth obtained the highest number of calls and, as a consequence, of false positives to filter followed by VarGenius-HZD and XHMM. After filtering we have selected candidate HDs to inspect in IGV according to disease gene and its association with the patient phenotype (in Supplementary Table 7 the sum of VisButNotFit and FitButNotVis) and obtained: 8 HDs from ExomeDepth; 5 from XHMM, 10 from VarGenius-HZD; 7 from HMZDelFinder. Only 6 events in total passed all the evaluation filters and 5 of them were confirmed through PCR (Table 6). VarGenius-HZD identified all these events and HMZDelFinder found 4 out of 5. We went back to unfiltered results from the other tools to see at which stage they were lost: XHMM did not find any of these HDs; ExomeDepth detected all of them but they were initially filtered out because of their low BF value (<5) (Table 2 and 3). Indeed, reducing the BF threshold in ExomeDepth increases the number of calls to assess. The excellent performance of VarGenius-HZD was particularly striking as it obtained the highest number of true positives at a low cost of variants to manually inspect.

**Table 6.** VRCIRD cases resolved by detection of a HD.

| sample | gene | region | XHMM | ExomeDepth (BF) | VarGenius-HZD | HMZDelFinder |
| --- | --- | --- | --- | --- | --- | --- |
| ID_A739 | RAX2 | 19:3772155-377224 | NO | 7.4 | YES | YES |
| CREv1_A392 | RAX2 | 19:3771519-3772224 | NO | 11 | YES | YES |
| CREv1_A348 | RP2 | X:46719422-46719537 | NO | 7.8 | YES | YES |
| CREv1_ARRP129 | RP2 | X:46719424-46719537 | NO | 9 | YES | YES |
| ID_A860 | RPGR | X:38186587-38186793 | NO | 9 | YES | NO |

#### Experimental validation of detected HDs

The five identified HDs were validated by PCR and/or Sanger sequencing in the following patients: first, a HD of the first two exons of the *RAX2* gene (Van de Sompele et al., 2019) was identified in two, reportedly unrelated subjects (a female and a male), who had a clinical diagnosis of autosomal recessive retinitis pigmentosa (Figure 4). Both patients were born in a small village of ~1000 inhabitants in Campania (Italy). We believe that a founder effect within this small isolated community could account for the fact that they both carried the same deletion. Second, a hemizygous deletion in the *RP2* gene was identified in two young male subjects (17- and 21-year-old) who were first-degree cousins and diagnosed with X-linked early-onset retinitis pigmentosa (OMIM #300757, http://www.omim.org/entry/). Finally, the fifth case was a male patient carrying a hemizygous deletion of exon 1 of the *RPGR* gene and his clinical presentation was consistent with a diagnosis of an X-linked Retinitis Pigmentosa associated with mutations in this gene (OMIM #312610, http://www.omim.org/entry/).

**Figure 4:**
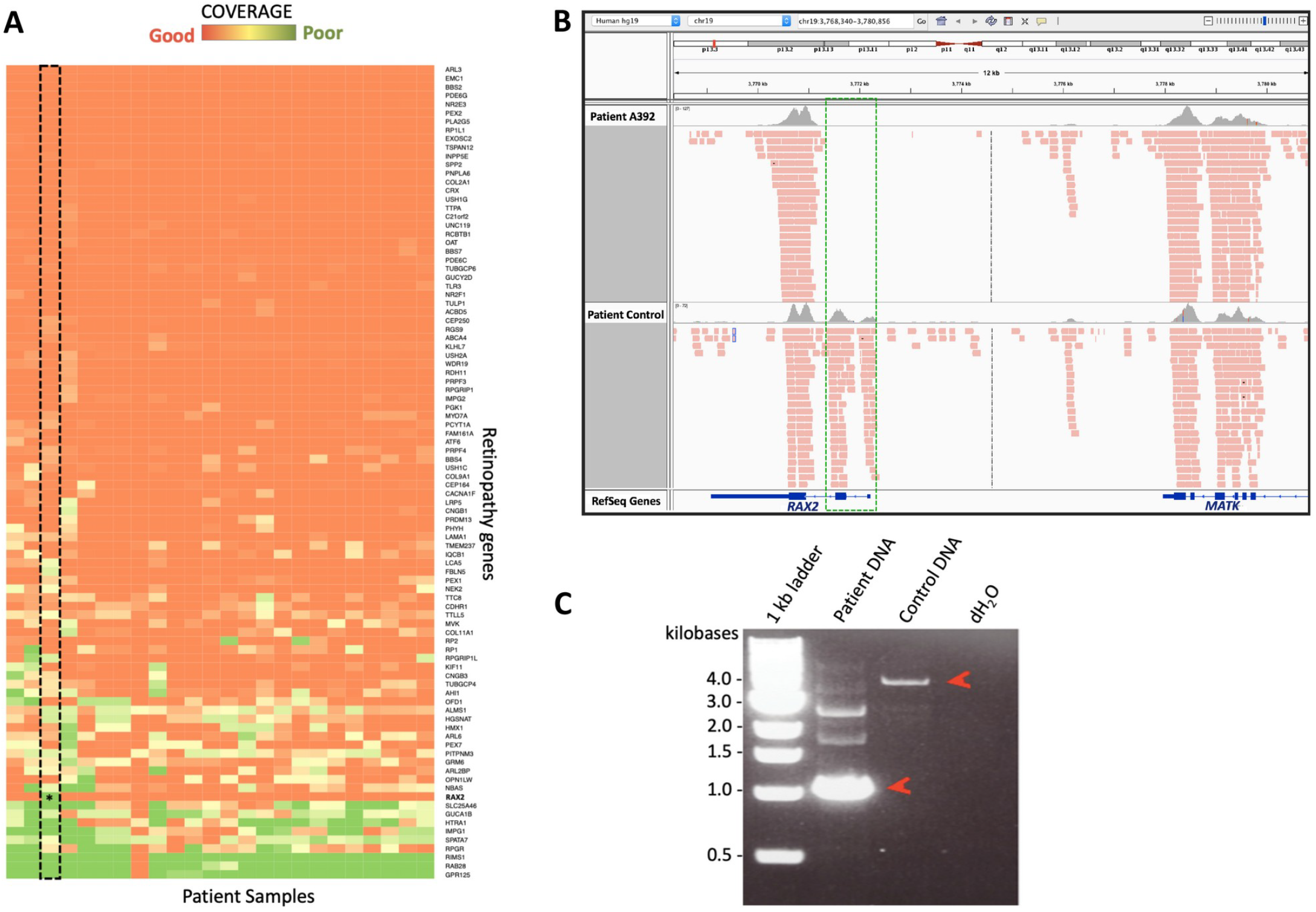
Experimental validation of the HD detected with our algorithm in the RAX2 gene. (A) Coverage heatmap of retinopathy genes in the WES data of IRD patients. Patient samples are shown in the x axis and gene names on the y axis. The extent of coverage is plotted according to the reported color scale. The RAX2 gene is well covered across all individuals, but poorly covered in A392 (asterisk in the framed column). (B) IGV coverage tracks for the alignment file from patient A392 (upper track) and a control patient (lower track). The lack of reads spanning the exon 2 of RAX2 in A392 (green box) suggested that the corresponding region was deleted in both alleles of the analysed proband. (C) PCR amplification of the genomic region spanning the identified deletion in the proband’s genomic DNA (‘Patient DNA’) and in a control DNA sample. The difference in size between the two amplicons (red arrowheads) indicated the presence of an extensive HD in the proband.

## DISCUSSION

HDs often lead to loss of function with pathogenic roles both in mendelian diseases and cancer (Cheng et al., 2017; Cox et al., 2005; Lupski et al., 2011; Valduga et al., 2015). Indeed, a significant percentage of human mendelian diseasesis are reported to be caused by molecular disruption within exons (Redon et al., 2006; Yuan et al., 2020). NGS based approaches became cheap during the last decade allowing diagnostic laboratories to use targeted sequencing (Levy & Myers, 2016). Nonetheless, the investigation of CNVs in WES is still challenging for several reasons (Guo et al., 2013; P. S. Samarakoon et al., 2016; Zare et al., 2017). State-of-the-art tools require as input several samples for such comparison that should be unrelated and sequenced with the same target (Fromer & Purcell, 2014; Gambin et al., 2017; Plagnol et al., 2012). Yet, comparative works have demonstrated a high number of false positives and hence alternative CNV detection strategies and filtering methods are needed (Feng et al., 2015a; Guo et al., 2013; Tan et al., 2014).

The goal of this work was to explore different solutions for HDs discovery in targeted sequencing and to automate the overall workflow. We developed VarGenius-HZD that searches for HDs within the single sample and leverages multi-sample information to corroborate such calls and we integrated it within our recently developed VarGenius. Being able to automate the CNV analysis workflow executing state-of-the-art tools, this software is extremely useful for those laboratories lacking bioinformatics expertise. To make VarGenius-HZD useful for researcher exploiting other softwares for the variant calling, we developed also a stand-alone VarGenius-HZD; in this version the user should provide the list of full paths to the BAM files and the target file. One limitation of the stand-alone tool (compared to the complete VarGenius software) is that it cannot provide parents coverage as annotation but only average across all samples used.

To compare our algorithm with state-of-the-art methods, we applied VarGenius-HZD, ExomeDepth, HMZDelFinder and DECoN to 50 samples from 1KGP. The highest number of TPs was achieved only with our algorithm, hence it is more sensitive than state-of-the-art tools demonstrating that BoC can be effectively used to detect such kind of variants. Furthermore, our tool was able to detect correctly all the synthetic HDs that we inserted within randomly chosen samples in the same dataset, achieving a sensitivity of 100% while the only comparable results were obtained with HMZDelFinder with a sensitivity of 80%. ExomeDepth and DECoN were not able to detect any of the simulated HDs. Our results are in agreement with other comparative studies which describe ExomeDepth’s ability to discover long CNVs covering large chromosomal regions while missing events that affect less than 3 exons. However, DECoN, which is based on ExomeDepth, provided similar results. We speculate that a higher number of TPs and thus higher sensitivity rather than precision would be preferable for clinical diagnosis at a cost of filtering few additional CNVs during downstream prioritization.

We then assessed the performance of VarGenius-HZD in a clinical context using targeted sequencing data from a cohort of unsolved IRD patients. Analysis of CNVs using ExomeDepth and XHMM with such data turned out to be challenging. These tools detect hundreds of events and filtering FPs was a tough task. We observed several false positives detected by ExomeDepth and XHMM, in agreement with current studies showing that state-of-the-art CNV calling algorithms are influenced by different instruments outcome and low coverage samples possibly due to the high number of off-target bases, duplicates and low base quality. We speculated that CNV callers should deal with such issues and, to reduce the false discovery rate, as a pre-processing step, it could be useful to remove outlier samples which have a high number of calls (e.g >2 standard deviation). After filtering we could confirm, through experimental assays, 5 pathogenic HDs. Only VarGenius-HZD was able to detect all of them. In summary: XHMM lost all of them; ExomeDepth detected all except one but provided very low BF score hence they were initially excluded; HMZDelFinder detected all except one. One of the called HDs was instrumental in defining a new association of biallelic variants in the *RAX2* gene with autosomal recessive Retinitis pigmentosa (Van de Sompele et al., 2019).

In summary, the use of targeted sequencing data for CNV discovery, as well as the automation of this process (which currently requires programming skills) are of great importance. Here we report an algorithm that could be useful to identify rare HDs demonstrating that BoC is a valuable feature for their detection. Given the extensive use of targeted sequencing as a first tier method for molecular genetic diagnosis, our work has a great importance for research and clinical practice.

