## Supplementary Material for "VarGenius-HZD allows accurate detection of rare homozygous or hemizygous deletions in targeted sequencing leveraging breadth of coverage"

Supplementary Table 1. HDs called in 1KGP data.
Supplementary Table 2. 1KGP HDs in VarGenius-HZD result.
Supplementary Table 3. 1KGP HDs in ExomeDepth result.
Supplementary Table 4. 1KGP HDs in HMZDelFinder result.
Supplementary Table 5. 1KGP HDs in DECoN result.
Supplementary Table 6. Single nucleotide homozygous variants found in 1KGP samples.
Supplementary Table 7. Summary statistics of HDs found and filtered in the VRCIRD cohort.
Supplementary Figure 1. IGV screenshot of a false positive detected in 1KGP data.
Supplementary Figure 2. Deletion of genes *CFHR1* and *CFHR3* in sample NA20798 of 1KGP data.
Supplementary Figure 3. HD of gene *UGT2B28* in NA18504 1KGP data.

Supplementary Table 1. HDs called in 1KGP data.

| sample | chr | position | GT | Notes |
|---|---|---|---|---|
| NA19057 | chr12 | 10581591 | 1\|1:2.000:78,24,0 | VGII, ED(filt) |
| NA19137 | chr16 | 223447 | 1\|1:2.000:133,46,0 | VGII (filt), DECoN (filt BF:4.95) |
| NA18504 | chr4 | 70139616 | 1\|1:1.750:81,13,0 | VGII, ED (filt) |
| NA11919 | chr5 | 140222138 | 1\|1:2.000:573,259,0 | VGII,HMZ,ED,DECoN |
| NA19473 | chr9 | 107366951 | 1\|1:2.000:224,63,0 | Putative False Positive |

Supplementary Table 2. 1KGP HDs in VarGenius-HZD result.

| sample | BoC | DoC | AvgDoC | chr | start | end | gene | 1KGP |
|---|---|---|---|---|---|---|---|---|
| NA11919 | 0.17 | 8 | 3214.84 | 5 | 140235633 | 140238168 | PCDHAx | TP |
| NA11919 | 0.18 | 8 | 3208.58 | 5 | 140235633 | 140238021 | PCDHAx | TP |
| NA11919 | 0.18 | 6 | 2719.3 | 5 | 140235633 | 140237232 | PCDHAx | TP |
| NA20798 | 0 | 0 | 178.8 | 1 | 196795958 | 196796135 | CFHRx | NewTP |
| NA20798 | 0 | 0 | 162.88 | 1 | 196748291 | 196748486 | CFHRx | NewTP |
| NA20798 | 0 | 0 | 152.02 | 1 | 196748344 | 196748486 | CFHRx | NewTP |
| NA19057 | 0 | 0 | 52.76 | 12 | 10583715 | 10583827 | KLRCx | TP |
| NA18504 | 0.11 | 1 | 87.48 | 4 | 70146218 | 70146939 | UGT2B28 | NewTP |
| NA19137 | 0 | 0 | 20.72 | 19 | 55281302 | 55281336 | KIRx | TP(filt) |
| NA19137 | 0 | 0 | 38.74 | 16 | 223470 | 223599 | HBA2 | TP(filt) |

Supplementary Table 3. 1KGP HDs in ExomeDepth result.

| sample | nexons | CNV | BF | reads.expected | reads.observed | reads.ratio | 1KGP |
|---|---|---|---|---|---|---|---|
| NA20798 | 12 | 1:196744018-196801129 | 73.5 | 337 | 0 | 0 | NewTP |
| NA11919 | 1 | 5:147553798-147553933 | 16.4 | 76 | 0 | 0 | TP |
| NA19057 | 6 | 12:10583717-10588585 | 12.6 | 75 | 13 | 0.17 | TP |
| NA18504 | 6 | 4:70146220-70160527 | 22 | 103 | 2 | 0.02 | NewTP |

Supplementary Table 4. 1KGP HDs in HMZDelFinder result.

| sample | gene | CNV | 1KGP |
|---|---|---|---|
| NA11919 | PCDHA10 | 5:140235634_140236833 | TP |
| NA20798 | CFHR1 | 1:196795959_196796135 | NewTP |
| NA07347 | GHR | 5:42629139_42629205 | FP |
| NA12342 | GHR | 5:42629139_42629205 | FP |
| NA19213 | GHR | 5:42629139_42629205 | FP |
| NA18553 | CES1 | 16:55866915_55866967 | FP |
| NA18856 | ZNF630 | X:47918257-47919256 | FP |
| NA19137 | OR5P2 | 11:7817521_7818489 | FP |
| NA19236 | OR5P2 | 11:7817521_7818489 | FP |

Supplementary Table 5. 1KGP HDs in DECoN result.

| sample | nexons | CNV | BF | Reads.expected | Reads.observed | Reads.ratio | 1KGP |
|---|---|---|---|---|---|---|---|
| NA19137 | 2 | 16:223125-223599 | 4.59 | 21 | 0 | 0 | TP(filt) |
| NA11919 | 7 | 5:140235635-140237232 | 287 | 1554 | 0 | 0 | TP |
| NA11919 | 1 | 5:147553798-147553933 | 16 | 75 | 0 | 0 | TP |
| NA20798 | 13 | 1:196744017-196801129 | 71.2 | 327 | 0 | 0 | NewTP |
| NA19057 | 6 | 12:10583717-10588585 | 15.8 | 90 | 13 | 0.14 | TP |
| NA18504 | 7 | 4:70146220-70160527 | 23.7 | 109 | 1 | 0.01 | NewTP |
| NA19473 | 3 | 9:107379530-107380128 | 21.9 | 143 | 4 | 0.03 | FP |

Supplementary Table 6. Single nucleotide homozygous variants found in 1KGP samples.

| sample | chr | position | ID | REF | ALT | GT | Notes |
|---|---|---|---|---|---|---|---|
| NA20798 | chr1 | 196733401 | esv2672010 | G | <DEL> | 1\|1:2.000:239,112,0 | Close HD detected: VG,HMZ,ED,DC |
| NA20798 | chr1 | 196772601 | esv2672625 | G | <DEL> | 1\|1:2.000:169,82,0 | Close HD detected: VG, HMZ,ED,DC |
| NA19137 | chr19 | 55225201 | esv2671585 | T | <DEL> | 1\|1:1.550:77,12,0 | |
| NA19137 | chr19 | 55280101 | esv2666421 | C | <DEL> | 1\|1:2.000:92,41,0 | |
| NA19137 | chr19 | 55281501 | esv2661467 | C | <DEL> | 1\|1:2.000:161,63,0 | |
| NA18504 | chr4 | 70124301 | esv2656947 | T | <DEL> | 1\|1:2.000:1495,707,0 | Close HD detected: VG, ED |
| NA18950 | chr6 | 29872907 | esv2659548 | A | <DEL> | 1\|1:2.000:282,60,0 | |

Supplementary Table 7. Summary statistics of HDs found and filtered in the VRCIRD cohort.

| | ED | | | XHMM | | | VarGenius-HZD | | | HMZDelFinder | | |
|---|---|---|---|---|---|---|---|---|---|---|---|---|
| | CREv1 | CCP | ID | CREv1 | CCP | ID | CREv1 | CCP | ID | CREv1 | CCP | ID |
| totResults | 1744 | 4542 | 13797 | 788 | 255 | 168 | 3100 | 8916 | 1162 | 70 | 923 | 1951 |
| totFiltered | 52 | 204 | 1118 | 22 | 18 | 24 | 74 | 745 | 84 | 12 | 76 | 146 |
| VisButNotFit | 0 | 0 | 0 | 1 | 0 | 0 | 3 | 5 | 1 | 3 | 3 | 1 |
| FitButNotVis | 6 | 2 | 0 | 4 | 0 | 0 | 0 | 0 | 1 | 0 | 0 | 0 |
| selPutativeHDs | 1 | 0 | 0 | 3 | 0 | 0 | 3 | 0 | 3 | 3 | 0 | 2 |
| physicianSelection | 1 | 0 | 0 | 0 | 0 | 0 | 3 | 0 | 2 | 3 | 0 | 1 |
| Validated-TP | 1 | 0 | 0 | 0 | 0 | 0 | 3 | 0 | 2 | 3 | 0 | 1 |
| Validated-FP | 0 | 0 | 0 | 0 | 0 | 0 | 0 | 0 | 0 | 0 | 0 | 0 |

Supplementary Figure 1. IGV screenshot of a false positive detected in 1KGP data in sample NA19473.

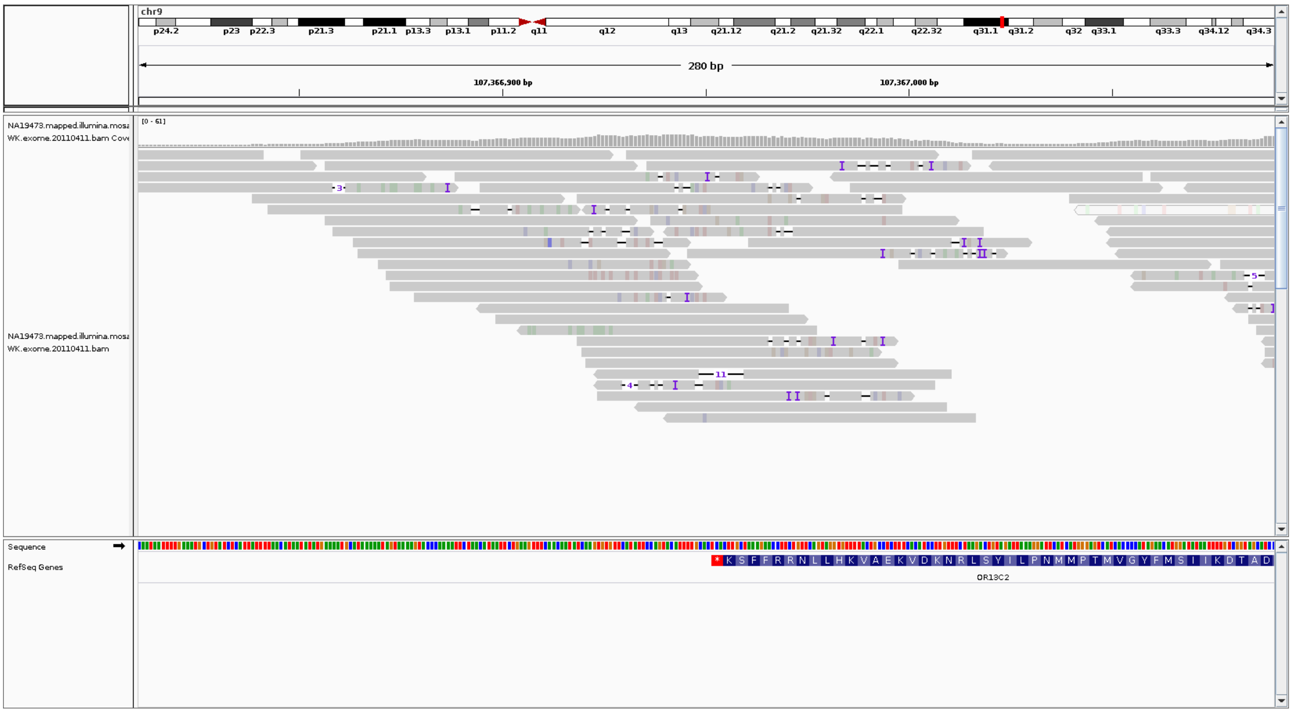

Supplementary Figure 2. Deletion of genes *CFHR1* and *CFHR3* in sample NA20798 of 1KGP data.

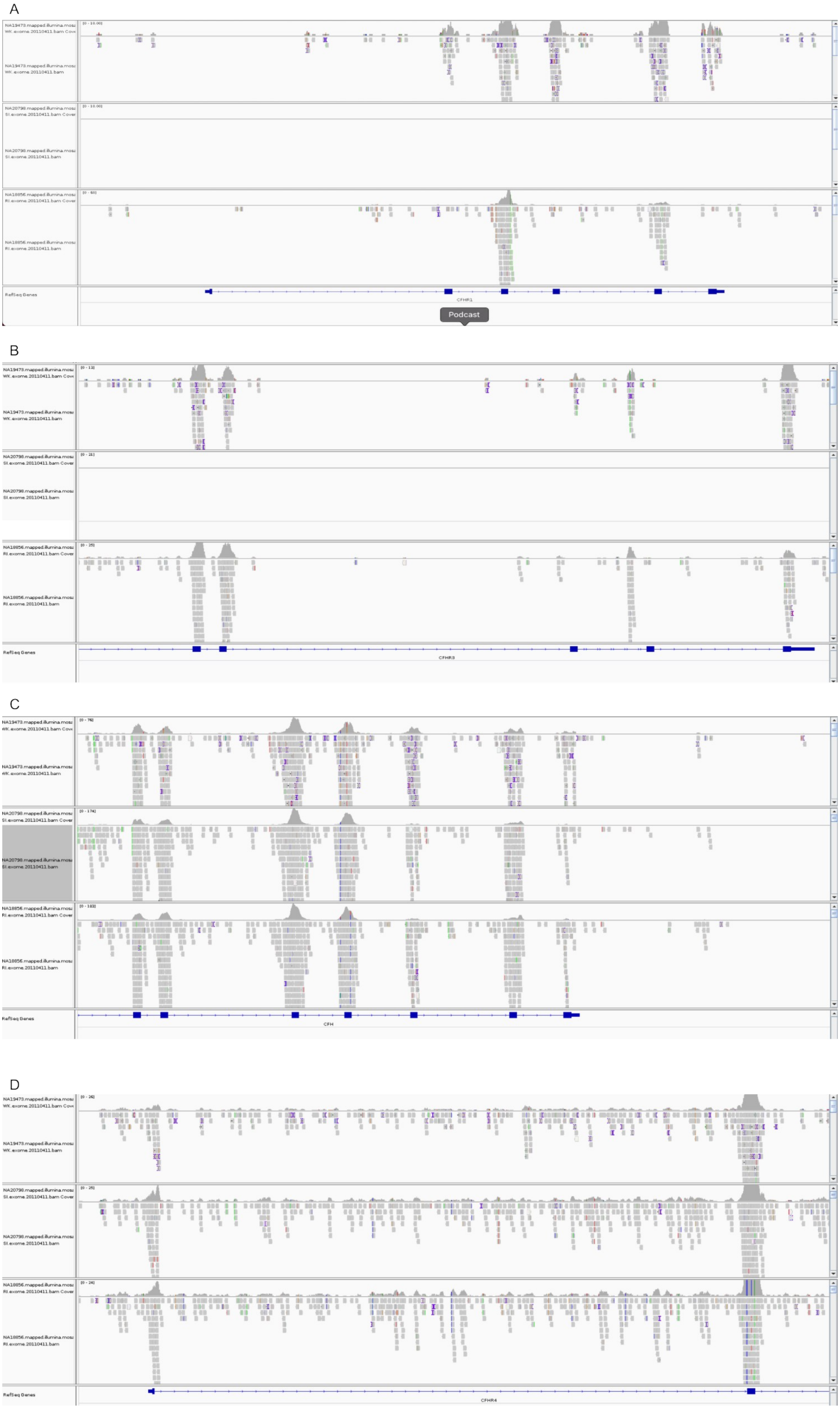

Supplementary Figure 3. HD of gene *UGT2B28* in NA18504 1KGP data.

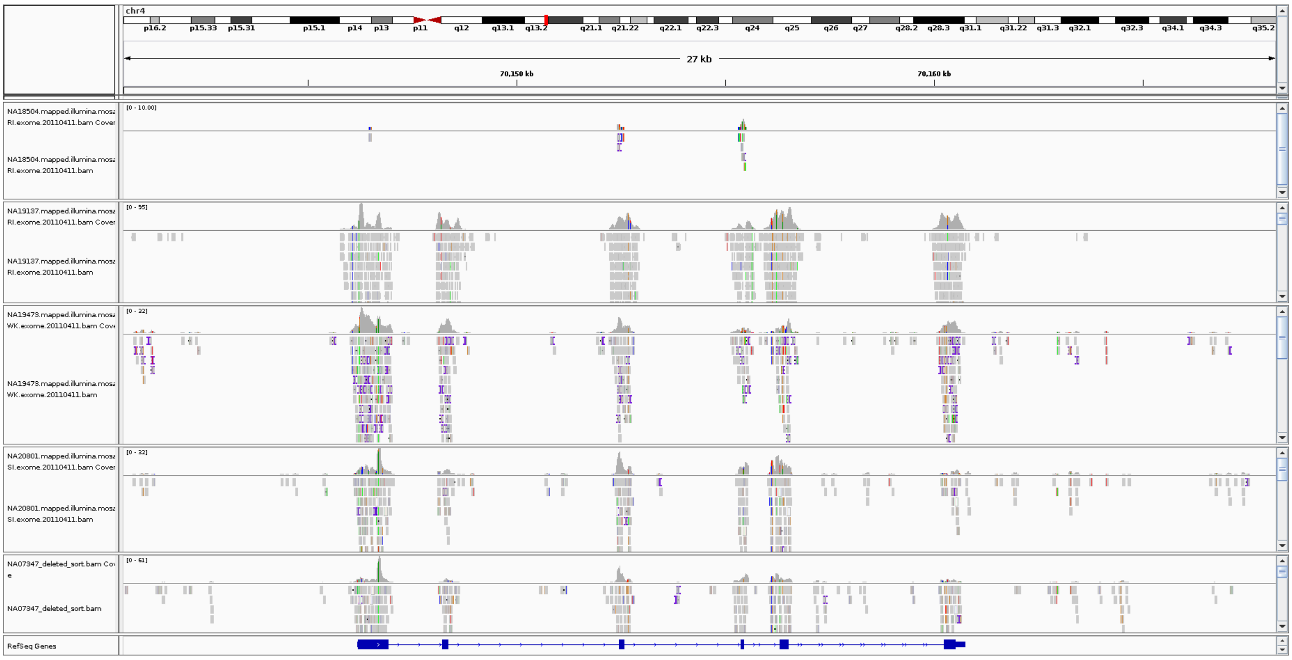
